# Actin Polymerization Status Regulates Tendon Homeostasis through Myocardin-Related Transcription Factor-A

**DOI:** 10.1101/2024.08.26.609684

**Authors:** Valerie C. West, Kaelyn Owen, Kameron L. Inguito, Karl Matthew M. Ebron, Tori Reiner, Chloe E. Mirack, Christian Le, Rita de Cassia Marqueti, Steven Snipes, Rouhollah Mousavizadeh, Dawn M. Elliott, Justin Parreno

## Abstract

The actin cytoskeleton is a potent regulator of tenocyte homeostasis. However, the mechanisms by which actin regulates tendon homeostasis are not entirely known. This study examined the regulation of tenocyte molecule expression by actin polymerization via the globular (G-) actin-binding transcription factor, myocardin-related transcription factor-a (MRTF). We determined that decreasing the proportion of G-actin in tenocytes by treatment with TGFβ1 increases nuclear MRTF. These alterations in actin polymerization and MRTF localization coincided with favorable alterations to tenocyte gene expression. In contrast, latrunculin A increases the proportion of G-actin in tenocytes and reduces nuclear MRTF, causing cells to acquire a tendinosis-like phenotype. To parse out the effects of F-actin depolymerization from regulation by MRTF, we treated tenocytes with cytochalasin D. Similar to latrunculin A treatment, exposure of cells to cytochalasin D increases the proportion of G-actin in tenocytes. However, unlike latrunculin A treatment, cytochalasin D increases nuclear MRTF. Compared to latrunculin A treatment, cytochalasin D led to opposing effects on the expression of a subset of genes. The differential regulation of genes by latrunculin A and cytochalasin D suggests that actin signals through MRTF to regulate a specific subset of genes. By targeting the deactivation of MRTF through the inhibitor CCG1423, we verify that MRTF regulates Type I Collagen, Tenascin C, Scleraxis, and α-smooth muscle actin in tenocytes. Actin polymerization status is a potent regulator of tenocyte homeostasis through the modulation of several downstream pathways, including MRTF. Understanding the regulation of tenocyte homeostasis by actin may lead to new therapeutic interventions against tendinopathies, such as tendinosis.

## Introduction

Reorganization of the actin cytoskeleton is a critical regulator of musculoskeletal cell behavior, and may mediate both biomechanical and biochemical signals in tissues such as tendons. Tendons are connective tissues that enable movement by passive force transfer from muscle onto bone. To allow force transfer and to resist high levels of tensile loading, tendons are rich in extracellular matrix, predominantly type I collagen (Col1) (Screen, Berk, Kadler, Ramirez, & Young, 2015). The resident fibroblastic tenocytes are elongated and have a specialized filamentous (F-)actin cytoskeleton that orients longitudinally along collagen fibers (Ralphs, Waggett, & Benjamin, 2002). Healthy tenocytes express alpha-smooth muscle actin (αsma), a highly contractile isoform of actin (Ippolito, Natali, Postacchini, Accinni, & De Martino, 1977; M. Spector, 2001). αsma is thought to enable cell shape recovery following tissue elongation. Thus, tendon matrix and cells are specially equipped to transmit and withstand mechanical loading (Dede Eren, Vermeulen, Schmitz, Foolen, & de Boer, 2023).

Tenocytes are mechanosensitive, and in response to changes in the mechanical environment, tenocytes can adapt by shifting tendon homeostasis. For instance, an increased mechanical load (physiological overloading) causes an anabolic shift in tendon homeostasis. Overloading releases active transforming growth factor beta (TGFβ) from the extracellular matrix (T. Maeda et al., 2011). The resultant increase in TGFβ signaling increases the expression of scleraxis (Scx), a transcription factor considered to be a master regulator of tenogenic differentiation. The upregulation of Scx elevates the expression of Col1, but also Tenascin-C (Tnc) levels (Chiovaro, Chiquet-Ehrismann, & Chiquet, 2015; Espira et al., 2009; Lejard et al., 2007), a glycoprotein that is necessary for proper extracellular matrix organization (Kannus, 2000; Mehr, Pardubsky, Martin, & Buckwalter, 2000). This anabolic response leads to tendon growth to withstand the elevated load. In contrast to physiological overloading, stress deprivation causes a catabolic shift in tendon homeostasis. Stress deprivation reduces the expression of Col1, Tnc, Scx, and αsma and increases in chondrogenic molecules (SRY-Box Transcription Factor; Sox9) and matrix degradative (matrix metalloproteinases; Mmps) expression (Arnoczky, Lavagnino, Egerbacher, Caballero, & Gardner, 2007; Egerbacher et al., 2022; Gardner, Arnoczky, Caballero, & Lavagnino, 2008; Inguito et al., 2022; Mousavizadeh, West, Inguito, Elliott, & Parreno, 2023; Wunderli et al., 2020). This catabolic shift in expression leads to tendon degeneration and is consistent with a tendinosis-like phenotype.

Reorganization of F-actin may play a critical role in regulating tendon homeostasis by tenocytes. In support of the critical role that F-actin may play, the catabolic shift in tenocytes coincides with a loss of F-actin (Inguito et al., 2022; Lavagnino & Arnoczky, 2005). Additionally, perturbation of actin networks in cultured tenogenic cells induces tendinosis-like molecular expression. In mesenchymal cells, exposure to latrunculin A to depolymerize actin results in the repression of tenogenic molecules, type I collagen (Col1), and scleraxis (Scx) (Maharam et al., 2015). Similarly, preventing myosin binding along F-actin by exposing cells to blebbistatin represses the tenogenic phenotype by reducing Col1 and Scx in mesenchymal cells (Maharam et al., 2015). Blebbistatin also lowers αsma and increases tenocyte expression of Mmp-1, −3, −14 (Jones et al., 2023; E. Maeda, Sugimoto, & Ohashi, 2013). Furthermore, we determined that inhibition of F-actin stabilization molecule, tropomyosin 3.1, leads to actin depolymerization resulting in the downregulation of tenogenic (Col1, Tnc, Scx, αsma) mRNA levels and increase in both chondrogenic (Acan, Sox9) and protease (Mmp-3, Mmp13) mRNA levels consistent with tendinosis gene changes (Inguito et al., 2022). While these studies highlight the importance of intact F-actin networks in tenocytes, the complete mechanisms by which dysregulation of F-actin networks promotes catabolic expression by tenocytes remains unclear.

Actin polymerization is known to regulate molecule expression through interactions with actin-binding transcription factors (Delve et al., 2020; Delve et al., 2018; Gonzalez-Nolde et al., 2024; Mgrditchian et al., 2023; Nalluri, O’Connor, & Gomez, 2015; Parreno et al., 2014). Myocardin-related transcription factor-a (MRTF) (also known as megakaryocyte leukemia factor-1; MKL1) is an actin-binding transcription factor known to regulate the expression of several molecules associated with healthy tenocytes. To positively regulate gene expression, MRTF interacts with serum response factor (SRF), which binds to a 10-bp CArG box on the promoter region of genes to enhance expression (Sun et al., 2006). However, MRTF also has a high affinity for monomeric globular (G-)actin (Mouilleron, Langer, Guettler, McDonald, & Treisman, 2011). An increase in G-actin, caused by actin depolymerization, increases the binding of MRTF to G-actin. The binding of MRTF to G-actin causes MRTF to be sequestered in the cytoplasm, leading to the downregulation of MRTF-regulated genes. In other cell types, MRTF inhibition downregulates the expression of matrix molecules, Col1 and Tnc (Asparuhova, Ferralli, Chiquet, & Chiquet-Ehrismann, 2011; Luchsinger, Patenaude, Smith, & Layne, 2011), and αsma (Parreno, Raju, Wu, & Kandel, 2017; Xie, Ning, Zhang, Ni, & Ye, 2022). Nevertheless, it remains unclear if MRTF regulates the expression of these genes in tenocytes. Furthermore, actin depolymerization in tenocytes has other profound effects, such as downregulation of Scx, upregulation of chondrogenic expression (Sox9), and upregulation of Mmps (Mmp3 and Mmp13) (Inguito et al., 2022). The regulation of these genes by actin polymerization through MRTF is yet unclear.

Understanding the role actin-based signaling plays in controlling tendon homeostasis through the regulation of tenocyte transcription will provide insight into tendon disease and repair. In tendinosis, abnormal biomechanical and biochemical signals may converge on the F-actin cytoskeleton to regulate tissue homeostasis. While the understanding of the regulation of actin by biomechanical forces has been investigated, the regulation of actin by biochemical mediators is less clear. While TGFβ is associated with pro-anabolic effects and is regarded as the critical pathway for mammalian tendon formation and regeneration (Havis et al., 2016; Kaji, Howell, Balic, Hubmacher, & Huang, 2020; Kuo, Petersen, & Tuan, 2008; Pryce et al., 2009), the regulation of actin by TGFβ in tenocytes is not known. In other cell types, TGFβ is a known regulator of F-actin. Since TGFβ is capable of enhancing Col1 synthesis (Heinemeier, Langberg, Olesen, & Kjaer, 2003), promoting the expression of Scx (Brown, Galassi, Stoppato, Schiele, & Kuo, 2015), and suppressing matrix Mmp activity (Farhat et al., 2015), an understanding on the actin-based regulation of molecular may provide new insights into the stimulating tendon anabolism and regeneration.

This study aims to examine the regulation of tenocyte phenotype by actin. We hypothesize that actin polymerization status is regulated biochemically by TGFβ and actin polymerization status regulates tendon homeostasis by mediating tenocyte gene expression partly through MRTF.

## Materials and Methods

### Mice and tendon cell isolation

Wild-type C57Bl6/J mice were obtained from The Jackson Laboratory (Bar Harbor, ME) and bred following approved animal protocols from the University of Delaware Institutional Animal Care and Use Committee (IACUC). Female and male mice aged 8-10 weeks were euthanized by CO2 inhalation and used for these studies. Tendon cells were isolated from both sexes. However, our studies indicated no sex differences in response to treatments. Therefore, data represents combined data obtained from experiments on cells from both sexes.

Tendon cells were isolated as previously described (Inguito et al., 2022). Briefly, tendon fascicles were dissected from tails and placed in Dulbecco’s Modified Eagles’ Media (DMEM), consisting of 1% antimycotic/antibiotic. After 15 minutes, fascicles were transferred into 0.2% collagenase A (MilliporeSigma) and maintained at 37°C. Following overnight collagenase digestion, the digests were strained through a 100µm filter (GenClone; Genesee Scientific; San Diego, CA, USA). Cells were then pelleted at 800g for 8 minutes. Pellets were resuspended in DMEM consisting of 10% fetal bovine serum (FBS; GenClone) and 1% antimycotic/antibiotic (defined as complete media). Cells were seeded at a density of ∼8.3 × 10^4^ cells/cm^2^ in either a six-well dish or on glass dishes. Media was replenished every three days with fresh, complete media.

### Treatment of cells with TGFβ1 or pharmacological inhibitors

Once tenocytes reached ∼50-70% confluency, complete media was replaced with DMEM media containing 0.5% FBS and 1% antimycotic/antibiotic (defined as serum starved media) +/− TGFβ1 or inhibitors. We used a concentration of 20ng/mL TGFβ1 (#7666-MB-005/CF; ThermoFisher Scientific; Waltham, MA, USA). The concentration of pharmacological agents used in this study was 2μM, 10uM, and 10uM for and Latrunculin A (#10010630; Cayman Chemical Company; Ann Arbor, MI, USA), Cytochalasin D (#11330; Cayman), and CCG1423 (#10010350; Cayman), respectively.

### Cell morphology and area analysis

Light microscopy images of cells on six-well dishes were captured using a Swiftcam camera (Swiftcam Technologies, Hong Kong) mounted on an Axiovert 25 inverted phase-contrast microscope (Zeiss, Jena, Germany) or a Zeiss Primovert Microscope (Zeiss). Cell morphology and area were then analyzed by tracing cells using FIJI software as previously described (Schofield et al., 2024). Circularity (C) was defined as C = 4π(cell area/cell perimeter^2^) ranging from 0 (elongated ellipse) to 1 (perfect circle). Cell area and circularity data from each set were combined and plotted using violin plots in Prism software (Graphpad; Boston, MA, USA).

### Confocal microscopy

Tendon cells on glass dishes were fixed by incubating cells in 4% paraformaldehyde at room temperature. After 15 minutes, cells were washed three times in phosphate-buffered saline (PBS; potassium phosphate, sodium chloride, sodium phosphate dibasic; GenClone). Cells were then permeabilized using permeabilization/blocking solution (3% Goat serum, 3% BSA, and 0.3% Triton) for 30 minutes.

To visualize G- and F-actin, tenocytes were stained with vitamin D binding protein conjugated to Alexa 488 (VitDBP-488; 1:100; RayBiotech; Peachtree Corners, GA, USA) and rhodamine-phalloidin (1:50; Biotium; Fremont, CA), respectively.

To visualize MRTF, tenocytes were incubated in permeabilization/blocking buffer consisting of anti-MRTF rabbit monoclonal IgG primary antibody (1:200; Cell Signaling Techology; Danvers, MA, USA). After an overnight incubation at 4°C, cells were washed in PBS, three times for 5 minutes per wash. Cells were then incubated in secondary antibody solution which contained anti-rabbit CF488 secondary antibody (1:100; Biotium), rhodamine phalloidin (1:50; Biotium), and Hoechst 33342 (1:500; Biotium). After a 1-hour incubation at room temperature, cells were washed in PBS three times for 5 minutes per wash.

Stained cells were coverslip mounted with antifade mounting medium (Drop-n-Stain Everbrite; Biotium) and then visualized using a Zeiss 880 microscope with a 40x objective (NA = 1.3 oil objective; z-stacks with a step size of 0.3μm). ZEN (Black Edition; Zeiss) was used to process images.

### Analysis and quantification of Fluorescent Microscopy Images

Ratiometric analysis of G/F-actin in cells was determined on maximum intensity projections of images on FIJI using a previously described protocol (Schofield et al., 2024). Cell boundaries were traced using phalloidin staining of F-actin as an indicator of cell borders. Fluorescence intensities were measured in each channel, and G/F-actin ratios were calculated by dividing the measured intensities of G-actin-stained VitDBP-488 by F-actin-stained rhodamine-phalloidin. The derived G/F-actin fluorescent intensities of cells were normalized to control averages within each set. Combined data from sets were used for statistical analysis and are presented in dot-plots showing averages ± standard deviation.

Ratiometric analysis of nuclear to cytoplasmic MRTF was performed on maximum intensity projections of confocal images using FIJI. Cell boundaries were traced using phalloidin staining of F-actin as an indicator of cell borders. Nuclear boundaries were traced using Hoechst staining as an indicator of nuclear borders. The nuclear to cytoplasmic ratio was calculated by dividing the mean fluorescent intensity of nuclei by the calculated mean fluorescent intensity of the cytoplasm. The derived nuclear-to-cytoplasmic ratios were normalized to control averages within each set. Combined data from sets were used for statistical analysis and are presented in dot-plots showing averages ± standard deviation.

### RNA isolation

Cells on six-well dishes were washed in PBS and then placed in TRIzol reagent (Sigma-Aldrich) to harvest RNA. RNA was isolated by phase separation in chloroform and then mixed with 100% ethanol. RNA purification was performed using RNA Clean and Concentrator-5 kit (Zymo Research; Irvine, CA, USA) as per the manufacturer’s instructions.

Relative real-time RT-PCR was performed on equal concentrations of cDNA using qPCRBio SyGreen Blue Mix (PCR Biosystems; London, UK) with previously validated primers (Inguito et al., 2022). The ΔΔCT method was used to calculate mRNA levels using 18S for normalization (Schmittgen & Livak, 2008). The derived mRNA levels were normalized to control averages within each set. Combined data from sets were used for statistical analysis and are presented in dot plots showing averages ± standard error.

### Triton fractionation and protein extraction

Triton insoluble and soluble fractions were extracted from cultured tenocytes as previously described (Schofield et al., 2024). Briefly, Triton soluble fractions (containing G-actin) were extracted from cells by incubating cells in cytoskeletal buffer (100mM NaCl, 3mM MgCl2, 300 mM Sucrose, 1mM EGTA, 10mM PIPES) consisting of 0.2% Triton at room temperature for 2 minutes. This Triton soluble fraction was collected, and radioimmunoprecipitation assay (RIPA) lysis concentrate (10x RIPA; Millipore Sigma) was added to each sample to obtain a final 1x RIPA buffer concentration. The insoluble fraction (containing F-actin) was collected by scraping the remaining cellular contents on the dish into cytoskeletal buffer consisting of 0.2% triton and 1x RIPA. Samples were kept at −80°C.

To extract total protein contents from cells, cells on dishes were briefly washed in PBS. Cells were then placed in 1xRIPA in PBS containing protease inhibitor and scraped from dishes. Total protein was quantified using a bicinchoninic acid (BCA) protein assay (Prometheus; Genesee Scientific).

### WES capillary electrophoresis

Protein quantification in samples were determined using WES capillary electrophoresis, as previously described (Parreno et al., 2022). For quantification of pan-actin in Triton fractionated samples, samples were first sonicated (Model Q55, Qsonica; Newton, CT, USA) and then equal volumes of Triton soluble and insoluble lysates were prepared according to the manufacturer’s instructions and separated using a 12-230 kDa Separation module kit (Protein Simple, San Jose, CA, USA). Actin in each sample was quantified by probing using a rabbit anti–pan-actin (1:100; #4968; Cell Signaling) antibody. Secondary labeling of proteins was performed using anti-rabbit HRP-conjugate (Protein Simple). The proportion of actin in the Triton-soluble portion was quantified by dividing the amount of Triton-soluble actin by total actin (sum of actin in Triton-soluble and -insoluble fractions). Combined data from sets were used for statistical analysis and are presented in dot plots showing averages ± standard deviation.

To determine COL1α1 and αSMA protein expression in protein lysates, 0.5μg/mL of sonicated lysates were loaded per well and separated using either a 12-230 kDa Separation module kit or a 66-440kDA Separation module kit and probed with an αSMA antibody (1:50, Abcam; ab7817) or COL1α1 antibody (1:200; Novus; NBP1-300543), respectively. Total protein values were determined using a Chemiluminescent WES Simple Western Size-Based Total Protein Assay (Protein Simple). The COL1α1 and αSMA protein levels were determined by normalization to the total protein amount for each sample. The data from individual sets were expressed as a percentage of control averages. Combined data from sets were used for statistical analysis and are presented in dot plots showing averages ± standard deviation.

### Statistical analysis

Experiments were replicated at least three times on separate occasions. Graphpad prism was used for statistical analysis. Pooled data from sets were examined for outliers using the ROUT method, which uses robust nonlinear regression and identifies outliers from nonlinear curve fits with reasonable power and few false positives. The maximum false discover rate was set at 1% (Motulsky & Brown, 2006). Unpaired T-tests were used to detect differences between two groups of data.

## Results

### The anabolic affects of TGFβ on gene expression coincides with actin polymerization and nuclear MRTF

TGFβ is pro-anabolic and is critical for tendon formation as well as regeneration (Brown et al., 2015; Farhat et al., 2015; Havis et al., 2016; Heinemeier et al., 2003; Kaji et al., 2020; Kuo et al., 2008; T. Maeda et al., 2011; Pryce et al., 2009). In other cell types TGFβ has been shown to increase polymerization and nuclear import of MRTF causing alterations in gene expression (Crider, Risinger, Haaksma, Howard, & Tomasek, 2011; Gupta, Korol, & West-Mays, 2013; Johnson et al., 2014; Korol, Taiyab, & West-Mays, 2016; Kumawat et al., 2016; Speight, Kofler, Szaszi, & Kapus, 2016). Here, we test the hypothesis that TGFβ1 regulation of gene expression coincides with enhancement of F-actin polymerization and nuclear MRTF localization.

To confirm the gene regulatory effects of TGFβ1, we exposed isolated tenocytes to TGFβ1 and performed real-time RT-PCR. After one day of treatment, we found that TGFβ1 upregulates tenogenic molecules (Col1, Tnc, Scx) and contractile isoform of actin, αsma (Figure 1). TGFβ1 did not alter Acan mRNA levels; however, TGFβ1 reduces Sox9 mRNA levels. Furthermore, TGFβ1 reduces Mmp3 and Mmp13 mRNA levels.

**Figure 1.**
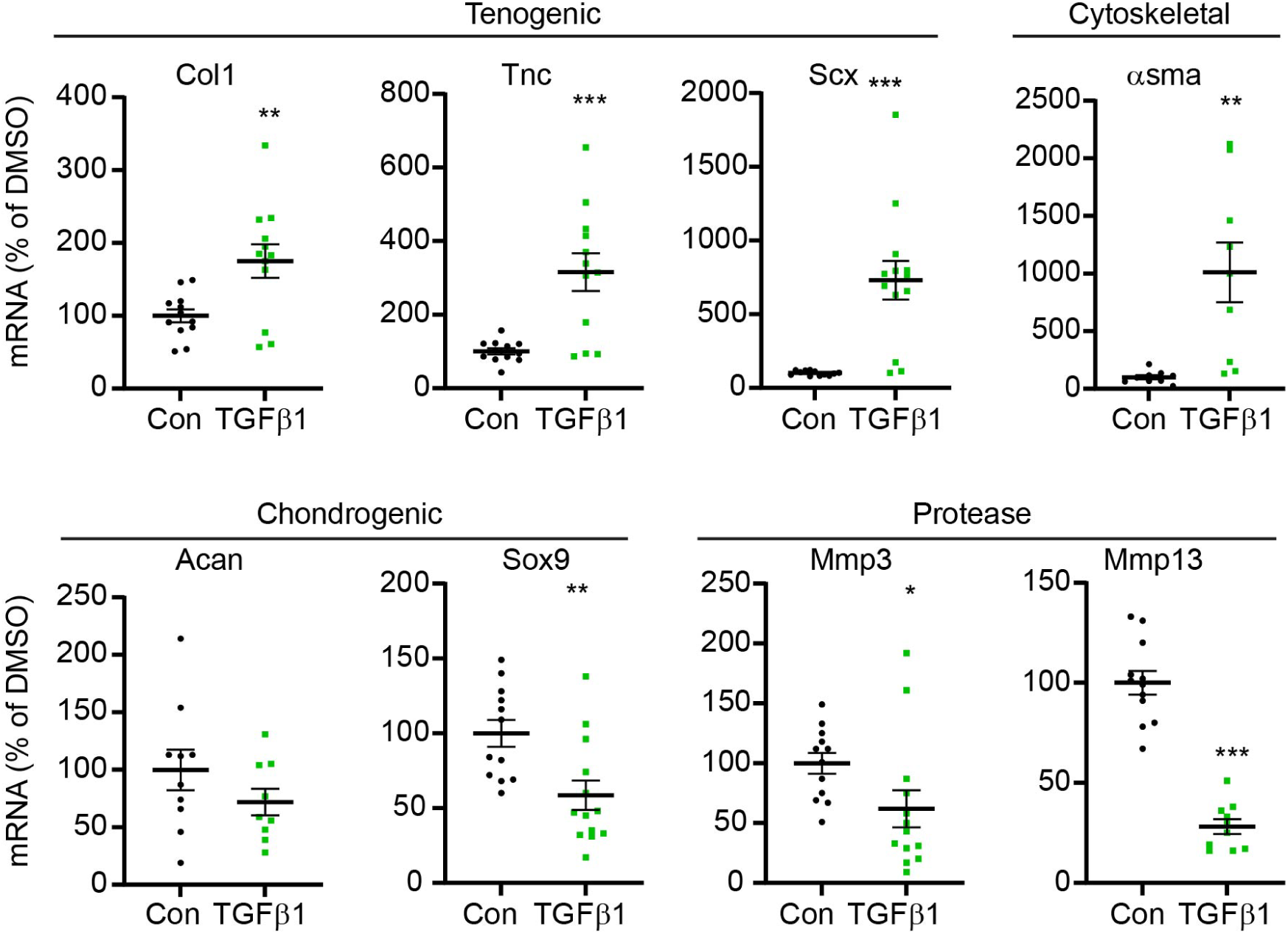
TGFβ1 treatment induces anabolic changes to tenocyte gene expression. Relative real-time PCR of tenocytes exposed to 20ng/mL TGFβ1 for 1day as compared to untreated control (Con) demonstrating an increase in tenogenic and asma mRNA levels, and a reduction in Sox9 as well as protease mRNA levels. Mean ± SEM is indicated on dot plots. *, p < 0.05; **, p < 0.01; ***, p < 0.001 as compared to Con.

In addition to the effects on gene expression, we also examined the impact of TGFβ1 on cell morphology (Figure 2A), actin organization (Figure 2B), and MRTF localization (Figure 2C). We found that TGFβ1 increases cell area (Figure 2A, D) and decreases circularity (Figure 2A, E). TGFβ1 also increases stress fibers in tenocytes (Figure 2B) and increases the proportion of F-actin in cells, as ratiometric analysis indicates that TGFβ1 decreases G/F-actin fluorescent staining intensity (Figure 2B, F).

**Figure 2.**
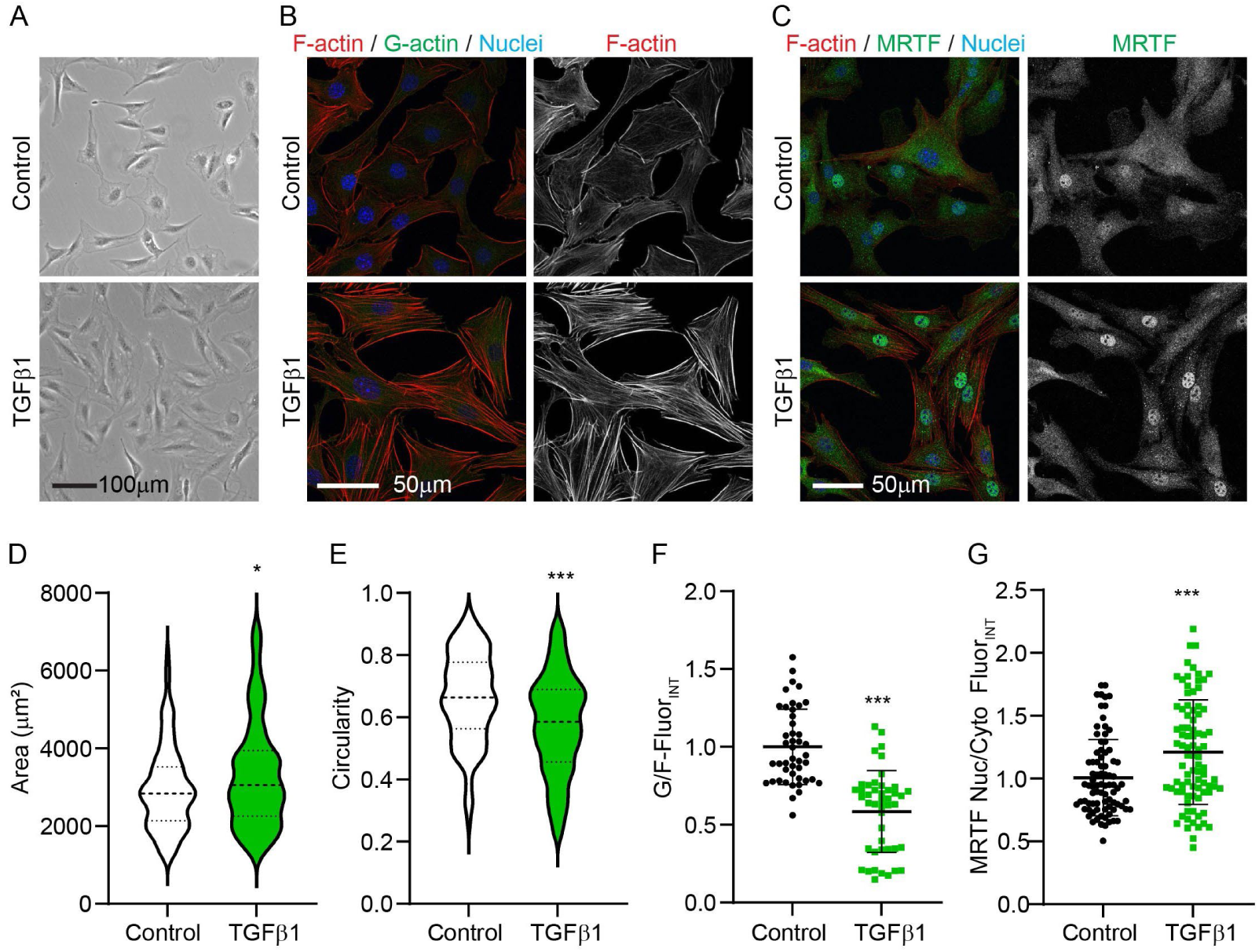
TGFβ1 alters cell morphology, actin organization/polymerization status, and MRTF localization. (A) Light and (B, C) confocal microscopy images of cells exposed to 20ng/mL TGFβ1 for 1 day. Quantification of light microscopy images in ‘A’ indicates, by violin plots, (D) an increase in cell area and a (E) reduction in circularity. Quantification of G-actin (Green; Vitamin D binding protein – Alexa 488) and F-actin (Red; Rhodamine-Phalloidin) fluorescence in ‘B’ demonstrates a (F) decrease in G/F-actin fluorescence. Quantification of MRTF (Green) nuclear and cytoplasmic mean fluorescence intensity in ‘C’ demonstrates an (G) increase in the proportion of nuclear MRTF. Cells are counterstained with Hoechst for visualization of Nuclei. *, p < 0.05; ***, p < 0.001 as compared to Control.

G-actin binds MRTF and sequesters it in the cytoplasm of cells (Mouilleron et al., 2011). We performed image quantification to determine the relative proportion of MRTF within the nucleus and cytoplasm of cells treated with TGFβ1 and determined that it enhances the proportion of nuclear MRTF in tenocytes (Figure 2C, G).

Overall, TGFβ1 treatment results in an anabolic shift in the tenocyte phenotype. The coinciding increase in the proportion of F-actin and nuclear MRTF leads us to the hypothesis that actin polymerization regulates tenocyte phenotype through MRTF.

### Latrunculin A abrogates stress fibers and increases G/F-actin and MRTF, leading to unfavorable alterations in tenocyte homeostasis

To determine the effect of F-actin depolymerization on tenocytes, we directly perturbed actin in isolated tenocytes by exposing cells to latrunculin A. Latrunculin A binds to monomeric G-actin preventing polymerization into F-actin (Coue, Brenner, Spector, & Korn, 1987; I. Spector, Shochet, Kashman, & Groweiss, 1983). In other cells, treatment with latrunculin causes cell rounding, abrogates stress fibers, increases G/F-actin, and reduces nuclear MRTF (Kuwahara, Barrientos, Pipes, Li, & Olson, 2005; Miralles, Posern, Zaromytidou, & Treisman, 2003; Parreno et al., 2014; Parreno et al., 2017).

We found that latrunculin A treatment alters tenocyte morphology by reducing cell area (Figure 3A, B) and increasing circularity (Figure 3A, C), leading to round cells. We confirm that latrunculin A abrogates stress fiber organization in isolated tenocytes, causing an increase in G/F-actin (Figure 3D, E).

**Figure 3.**
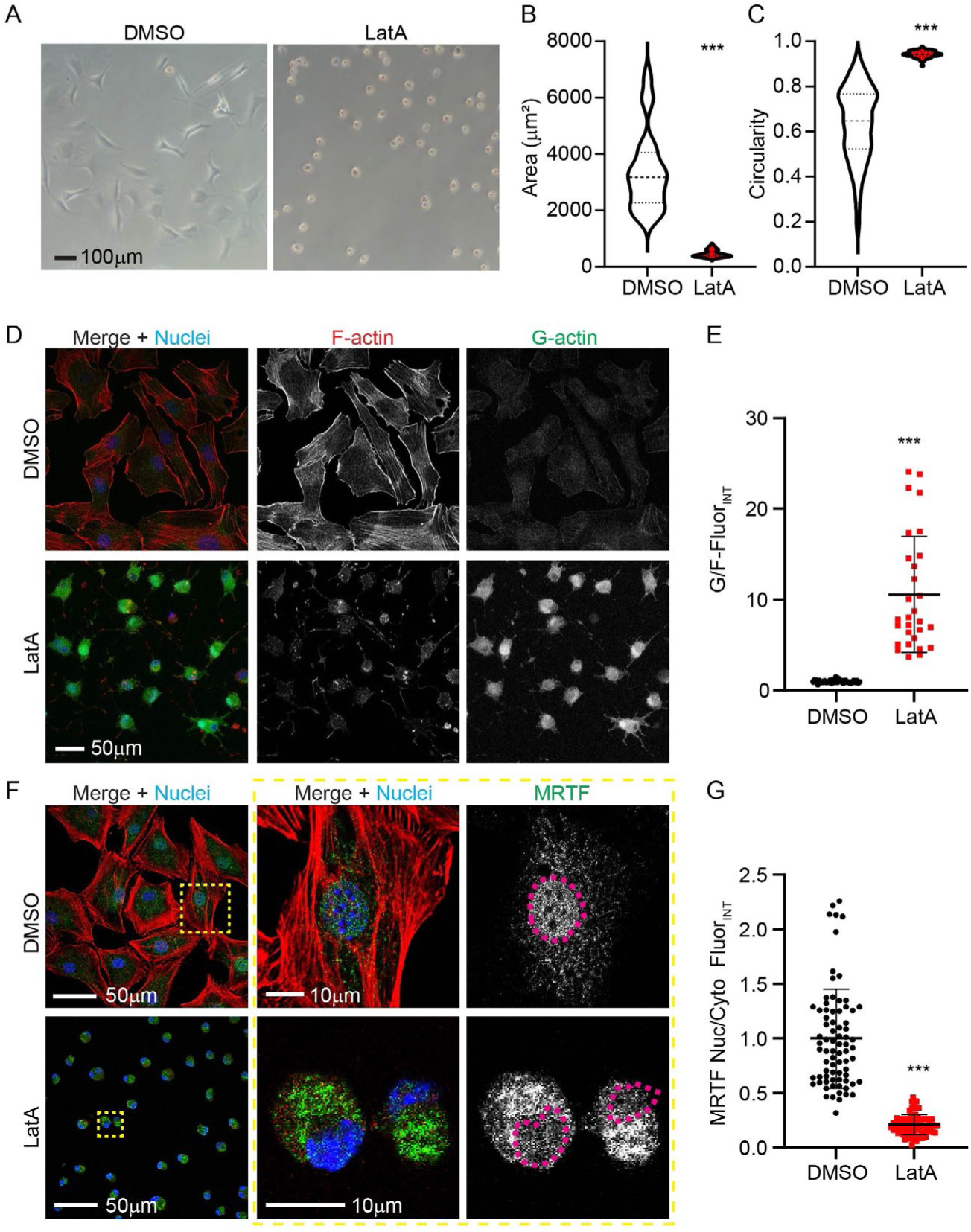
Latrunculin A induces cell rounding, an increase in G/F-actin, and a reduction in nuclear MRTF. (A) Light microcopy images and quantification, as shown by violin plots, for (B) area and (C) circularity of cells treated with 2μM latrunculin A for 1 day; latrunculin A induced cell rounding. (D) Confocal microscopy images of cells stained for F-actin (Red; Rhodamine-Phalloidin), G-actin (Green; Vitamin D binding protein – Alexa 488), and nuclei (Blue; Hoechst). (E) Quantification of G- and F-actin fluorescence demonstrates an increase in the proportion of G-actin. (F) Confocal microscopy images of cells stained for F-actin (Red; Rhodamine-Phalloidin), MRTF (Green), and Nuclei (Blue; Hoechst). Middle and right panels are zoomed in images of region marked by dotted yellow box in ‘F’. Right panel shows separated MRTF staining in gray, pink dash outlines represent nuclear borders. (G) Quantification of MRTF nuclear and cytoplasmic mean fluorescence intensity demonstrates a decrease in the proportion of nuclear MRTF. ***, p < 0.001 as compared to DMSO.

We next sought to determine if latrunculin A-induced actin depolymerization causes a reduction in nuclear localization of the G-actin binding transcription factor, MRTF. By staining for MRTF in fixed cells, we determined that latrunculin A reduces the proportion of nuclear MRTF (Figure 3F, G).

Finally, we examined the effect of Latrunculin A on tenocyte mRNA levels. Latrunculin A reduces Col1, Tnc, Scx, and αsma mRNA levels (Figure 4). Exposure of tenocytes to latrunculin A does not affect the mRNA levels of the cartilage matrix gene, Acan. However, latrunculin A increases the expression of the chondrogenic transcription factor, Sox9. Furthermore, latrunculin A treatment increases Mmp-3 and Mmp-13 mRNA levels.

**Figure 4.**
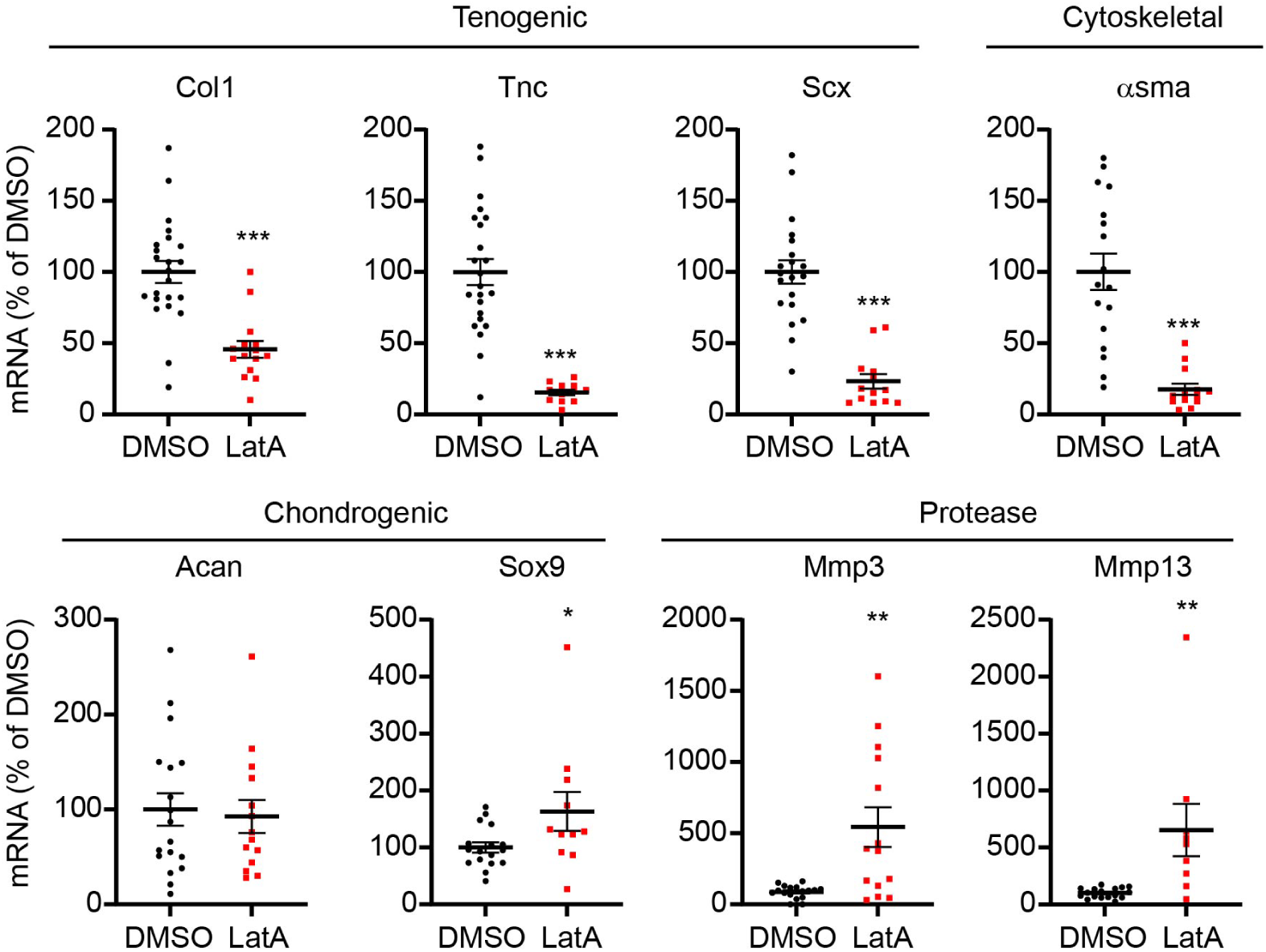
Latrunculin A treatment induces tendinosis-like gene expression changes. Relative real-time PCR of tenocytes exposed to 2μM latrunculin for 1day as compared to DMSO treated vehicle control demonstrating a reduction in tenogenic and αsma mRNA levels, and an increase in Sox9 as well as protease mRNA levels. Mean ± SEM is indicated on dot plots. *, p < 0.05; **, p < 0.01; ***, p < 0.001 as compared to DMSO.

Overall, the effects of latrunculin A treatment contrast the effects of TGFβ1 (Figure 1 and 2), promoting a catabolic shift in tenocyte phenotype. The coinciding increase in cytoplasmic MRTF localization by latrunculin-induced actin depolymerization led us to ask: does MRTF regulate gene expression in tenocytes?

### Cytochalasin D represses stress fibers, causing an increase in G/F-actin but also an increase in nuclear MRTF

To further elucidate the regulation of tenocyte phenotype by actin depolymerization and MRTF, we next exposed tenocytes to cytochalasin D, which also depolymerizes actin. The mechanism of action of cytochalasin D differs from that of latrunculin. Cytochalasin D attaches to the barbed end of F-actin, preventing barbed end monomer addition, whereas latrunculin sequesters G-actin to prevent polymerization (Carlier, Criquet, Pantaloni, & Korn, 1986; Cooper, 1987). First, we examined the effect of cytochalasin D on cell morphology (Figure 5A) and actin organization (Figure 5B-D). Similar to latrunculin treatment (Figure 3A-C), cytochalasin D decreases cell area (Figure 5A, E) and increases circularity leading to cellular rounding (Figure 5A, F). Cytochalasin D abrogates stress fibers (Figure 5B) and increases G/F actin fluorescence (Figure 5B, G). We determined that cytochalasin D treatment increases the proportion of actin in the Triton-soluble versus Triton-insoluble fraction of cells (Figure 5C, D, H). The increase in actin in the Triton-soluble fraction is consistent with an increase in G-actin within the cytoplasm of cells.

**Figure 5.**
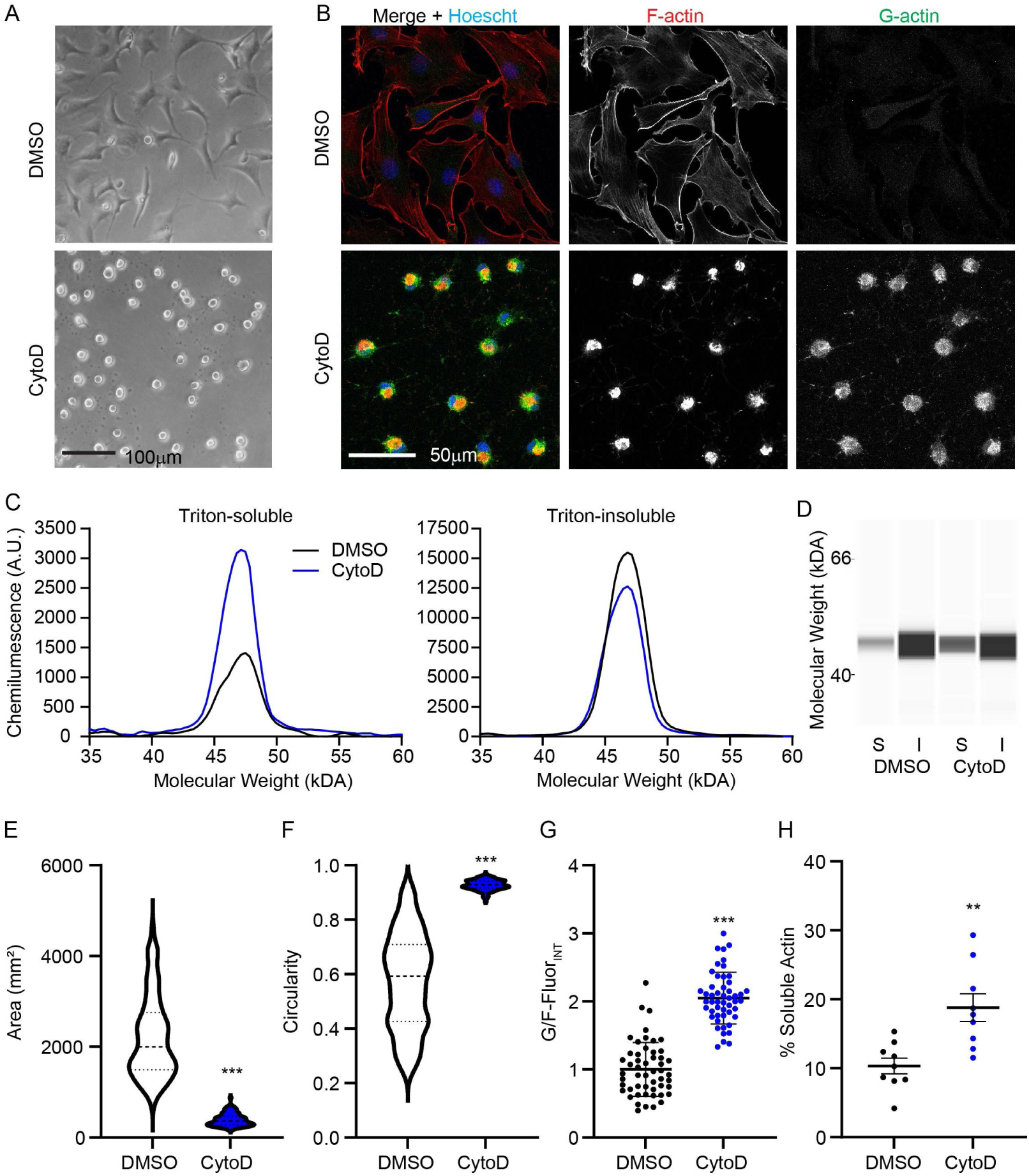
Cytochalasin D induces cell rounding, an increase in G/F-actin, and an increase in nuclear MRTF. (A) Light and (B) confocal microscopy images as well as capillary electrophoresis (C) spectropherograms and (D) pseudoblots for actin in triton-soluble ‘*S*’ and insoluble ‘*I*’ fractions of cells exposed to 10μM cytochalasin D. Quantification of (E) area and (F) circularity for cells treated with cytochalasin D for 1 day, by violin plots, in ‘A’ demonstrates that cytochalasin D induces cell rounding. (D, G) Quantification of F-actin (Red; Rhodamine-Phalloidin) and G-actin (Green; Vitamin D binding protein – Alexa 488) fluorescence in ‘B’ demonstrates that 2 hours of exposure to cytochalasin D increases in the proportion of G-actin. (H) Dot-plot of WES Capillary electrophoresis collective data as exemplified by representative findings in ‘C’ and ‘D’ demonstrate an increase in the proportion of actin in the soluble fraction of cells by 2 hours of exposure to cytochalasin D. **, p < 0.01; ***, p < 0.001 as compared to DMSO.

In contrast to latrunculin, which causes nuclear export of MRTF to deactivate MRTF signaling, cytochalasin D interferes between the interaction of G-actin and MRTF, causing an increase in nuclear MRTF (Medjkane, Perez-Sanchez, Gaggioli, Sahai, & Treisman, 2009). We confirmed that in contrast to latrunculin A treatment (Figure 3F, G), cytochalasin D increases nuclear MRTF localization (Figure 6A, B). We used the differential effect of cytochalasin D and latrunculin on MRTF to elucidate specific gene regulation by MRTF. Since cytochalasin D activates MRTF, genes that MRTF highly regulates would be upregulated by cytochalasin D treatment despite an increase in G/F-actin. Like latrunculin A treatment (Figure 4), cytochalasin D reduces the Col1 mRNA levels (Figure 6C) and increases Mmp-3 and Mmp-13 mRNA levels. However, compared to latrunculin A treatment, which decreases Tnc, Scx, and αsma mRNA levels (Figure 4), cytochalasin D increases Tnc, Scx, and asma mRNA levels (Figure 6C). The contrasting effects of latrunculin and cytochalasin D on specific mRNA levels suggest that actin depolymerization regulates this subset of genes in a manner that is highly dependent on MRTF.

**Figure 6.**
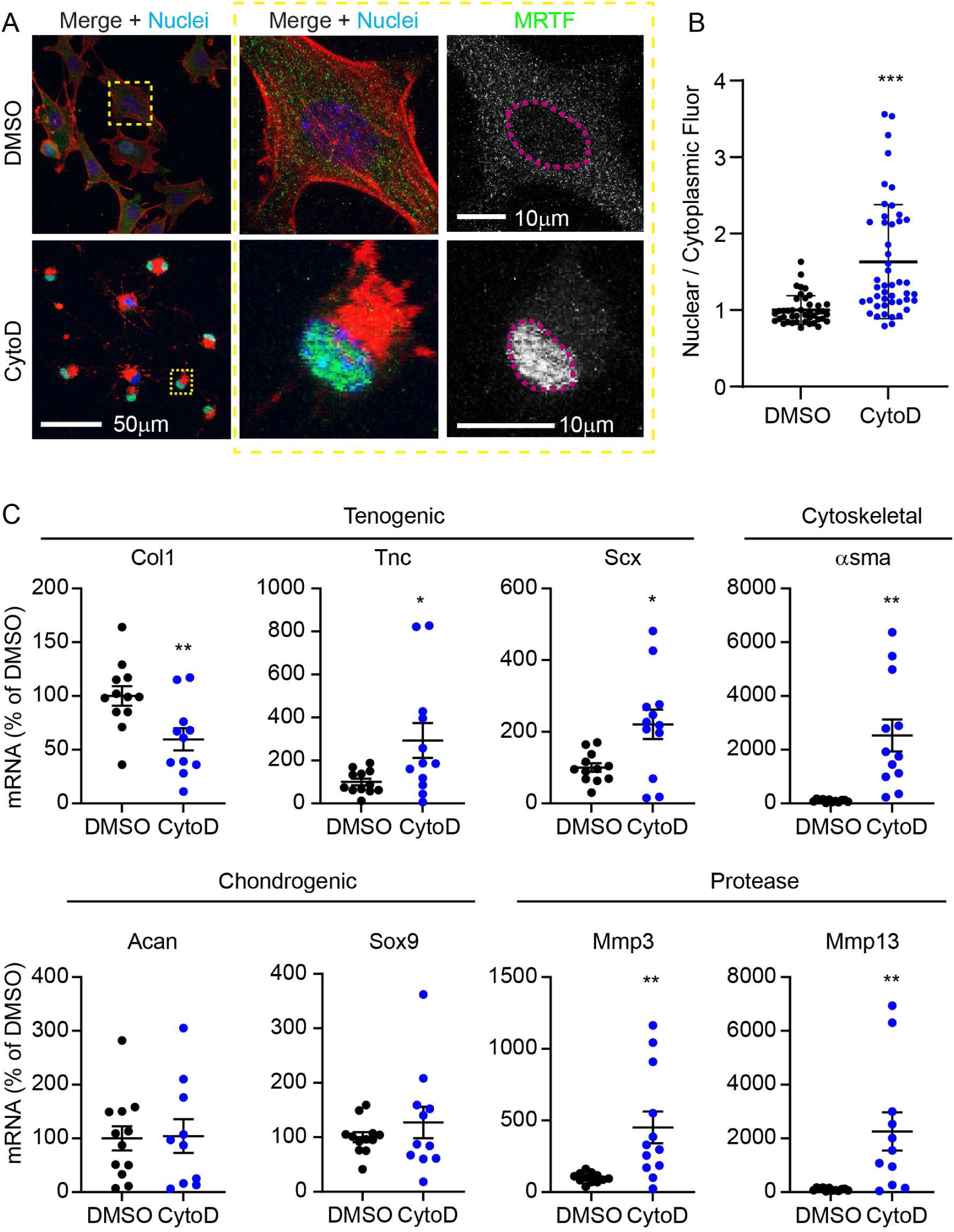
Cytochalasin D increases nuclear MRTF and differentially regulates gene expression. (A) Confocal microscopy images of cells treated with cytochalasin D for 4 hours. Cells were stained for F-actin (Red; Rhodamine-Phalloidin), MRTF (Green), and Nuclei (Blue; Hoechst). Middle and right panels are zoomed in images of region marked by dotted yellow box in left panel. Right panel shows separated MRTF channel in gray, pink dash outlines represent nuclear borders. (B) Quantification of MRTF nuclear and cytoplasmic mean fluorescence intensity demonstrates an increase in the proportion of nuclear MRTF. Relative real-time PCR of tenocytes exposed to 10μM cytochalasin D for 1day demonstrating altered regulation of genes, with opposite effects to latrunculin A treatment (Figure 4) for a subset of genes (Tnc, Scx, asma). Mean ± SEM is indicated on dot plots. *, p < 0.05; **, p < 0.01; ***, p < 0.001 as compared to DMSO.

### Pharmacological inhibition of MRTF reduces specific mRNA levels in tenocytes

To further elucidate the regulation of genes via MRTF, we exposed tenocytes to the MRTF inhibitor CCG1423. CCG1423 is an MRTF signaling deactivator that binds to the RPEL regions of MRTF-A and prevents nuclear import of MRTF (Hayashi, Watanabe, Nakagawa, Minami, & Morita, 2014). Compared to the actin depolymerization agents, latrunculin A or cytochalasin D treatment, CCG1423 had a more subtle effect on cell morphology. CCG1423 reduces the average cell area 1.8-fold, and cells became more elongated, with an average circularity of 0.50 (Figure 7A, E-F). In comparison, latrunculin A treatment reduces average cell area by 7.3-fold and causes cell rounding with an average circularity of 0.94 (Figure 3A-C). Furthermore, compared to latrunculin A treatment (Figure 3D), CCG1423 reduces but does not entirely abrogate, stress fibers. Furthermore, CCG1423 increased G/F-actin fluorescence intensity and soluble actin (Figure 7B-D, G-H). The degree of increase in G/F-actin fluorescence was much less than that of the latrunculin A treatment (1.3-fold versus 10.6-fold, respectively) (Figure 3D-E).

**Figure 7.**
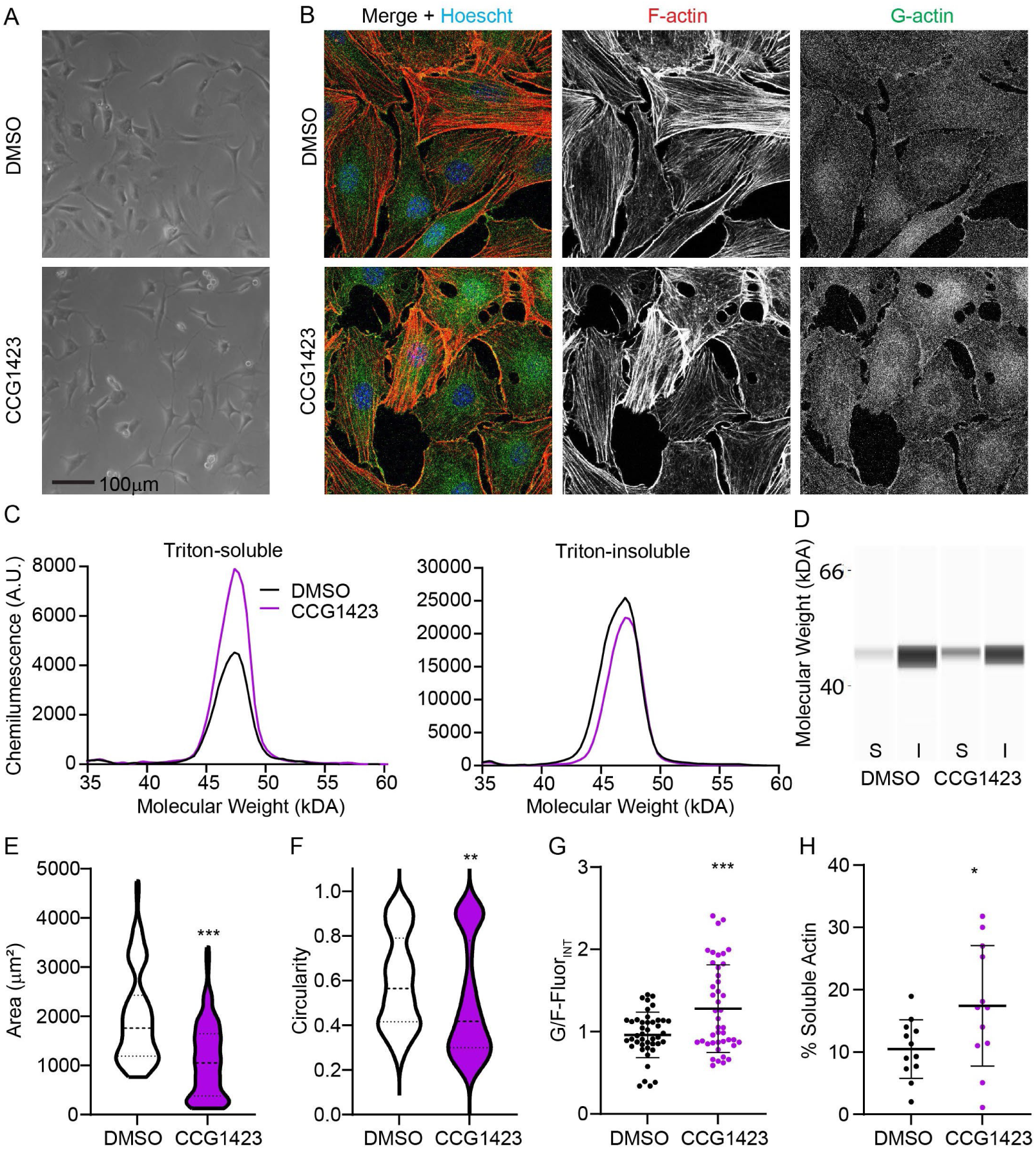
CCG1424 induces cell rounding, an increase in G/F-actin, and a decrease in nuclear MRTF. (A) Light and (B) confocal microscopy images as well as capillary electrophoresis (C) spectropherograms and (D) pseudoblots for actin in triton-soluble ‘*S*’ and insoluble ‘*I*’ fractions of cells exposed to 10μM CCG1423 for 1 day. Quantification of (E) area and (F) circularity for cells treated with CCG1423 in ‘A’ demonstrates cell rounding. (D, G) Quantification of F-actin (Red; Rhodamine-Phalloidin) and G-actin (Green; Vitamin D binding protein – Alexa 488) fluorescence in ‘B’ demonstrates that CCG1423 increases in the proportion of G-actin. (H) Dot-plot of WES Capillary electrophoresis collective data as exemplified by representative findings in ‘C’ and ‘D’ demonstrate an increase in the proportion of actin in the soluble fraction of cells by exposure of cells to CCG1423. *, p < 0.05; **, p < 0.01; ***, p < 0.001 as compared to DMSO.

CCG1423 treatment affected a subset, but not all, of the genes altered by latrunculin A. Similar to latrunculin A (Figure 3F, G), CCG1423 treatment reduces nuclear MRTF levels (Figure 8A, B). Also, in keeping with latrunculin A treatment, exposure of isolated tenocytes to CCG1423 reduces mRNA levels for Col1, Tnc, Scx, and αsma (Figure 8C). The effect of CCG1423 on Tnc, Scx, and αsma mRNA levels contrasts with the effects of cytochalasin D on these genes (Figure 6C). In contrast to latrunculin and cytochalasin D, CCG1423 did not affect Sox9, Mmp3, or Mmp13 mRNA levels (Figure 8C).

**Figure 8.**
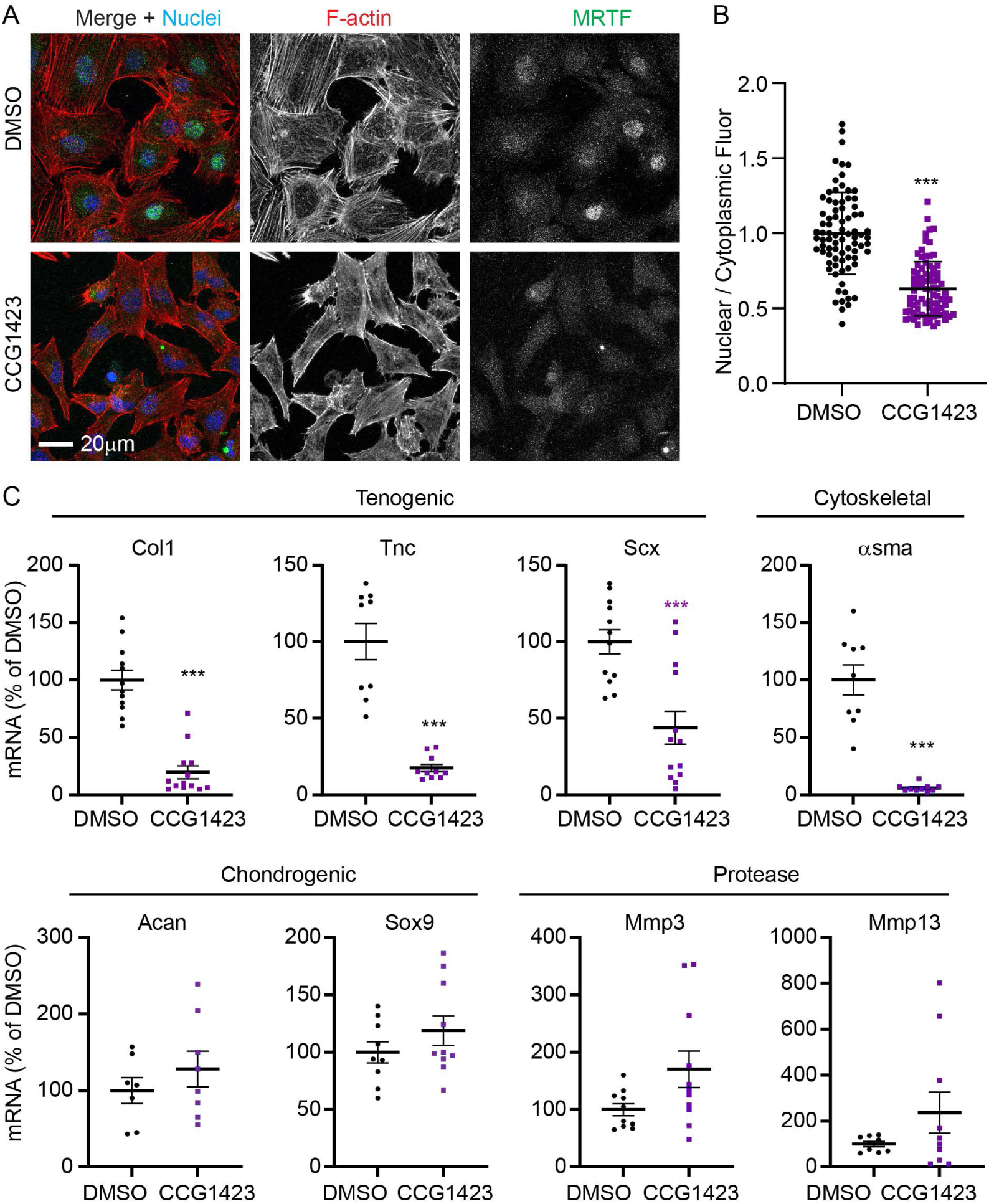
CCG1423 reduces nuclear MRTF and downregulates tenogenic and αsma mRNA levels. (A) Confocal microscopy images of cells treated with cytochalasin D for 1 day. Cells were stained for F-actin (Red; Rhodamine-Phalloidin), MRTF (Green), and Nuclei (Blue; Hoechst). (B) Quantification of MRTF nuclear and cytoplasmic mean fluorescence intensity demonstrates a decrease in the proportion of nuclear MRTF. (C) Relative real-time PCR of tenocytes exposed to CCG1423 demonstrates altered regulation of a subset of genes (Col1, Tnc, Scx, asma). Mean ± SEM is indicated on dot plots. *, p < 0.05; **, p < 0.01; ***, p < 0.001 as compared to DMSO.

Finally, we confirmed the effect of MRTF inhibition on select molecules at the protein level (Figure 9). Using WES capillary electrophoresis, we determined that CCG1423 reduces the expression of COL1 and aSMA protein. Collectively, our data shows that MRTF inhibition modestly affects cell morphology and F-actin. While alterations in molecular expression by CCG1423 treatment suggest a catabolic shift, MRTF regulates the expression of only a subset of actin-regulated molecules.

**Figure 9.**
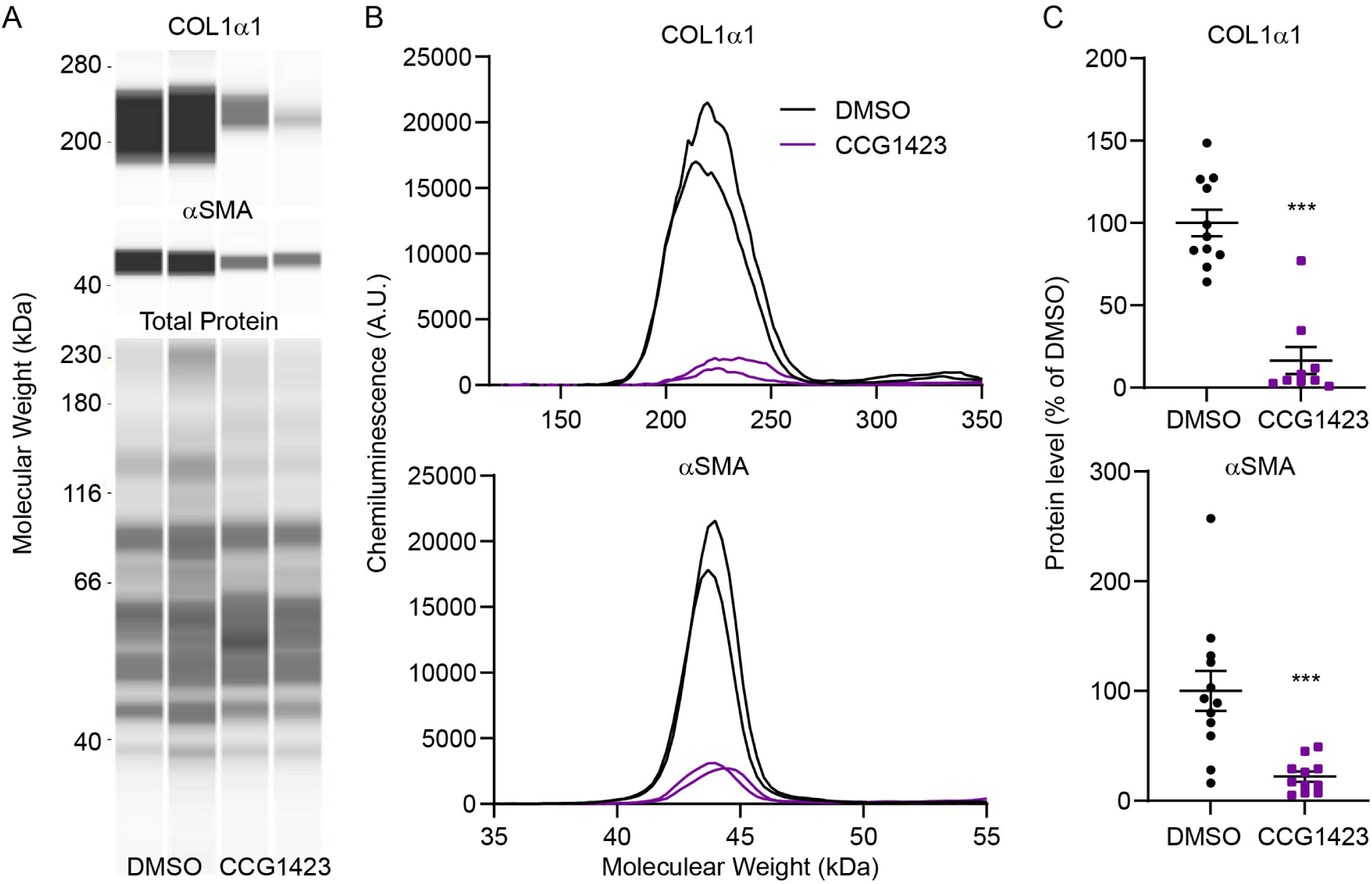
Modulation of select protein levels by exposure of tenocytes to 10μM CCG1423 for 2 days of treatment. Representative WES capillary electrophoresis. (A) pseudoblots and (B) electropherograms for COL1α1 and αSMA. (C) Dot-plots of COL1α1 and αSMA protein levels after normalization to total protein (example of total protein pseudoblot shown in ‘A’). ***, p < 0.001 as compared to DMSO treated cells.

## Discussion

This study reveals new mechanistic insights into regulating tenocyte homeostasis by actin. Our data supports the hypothesis that TGFβ1 can regulate F-actin, and that alterations in actin polymerization can regulate tenocyte molecule expression partly through MRTF (Figure 10). We found evidence that positive alterations to the tenocyte phenotype, through treatment with TGFβ1, coincide with elevated proportions of F-actin and nuclear MRTF. Therefore, we elucidated the regulation of tenocyte phenotype by actin through direct actin perturbation using latrunculin A as well as cytochalasin D. While latrunculin A and cytochalasin D treatment led to similar effects on morphology and actin organization, the varying effects on mRNA level regulation by the treatments could be attributed to differential regulation of MRTF. We confirmed the regulation of tenocyte phenotype by MRTF using CCG1423. CCG1423 had more subtle effects on morphology and F-actin than latrunculin A and downregulated a subset of genes. Thus, actin polymerization regulates tenocyte homeostasis in both an MRTF-independent and dependent manner (Figure 10).

**Figure 10.**
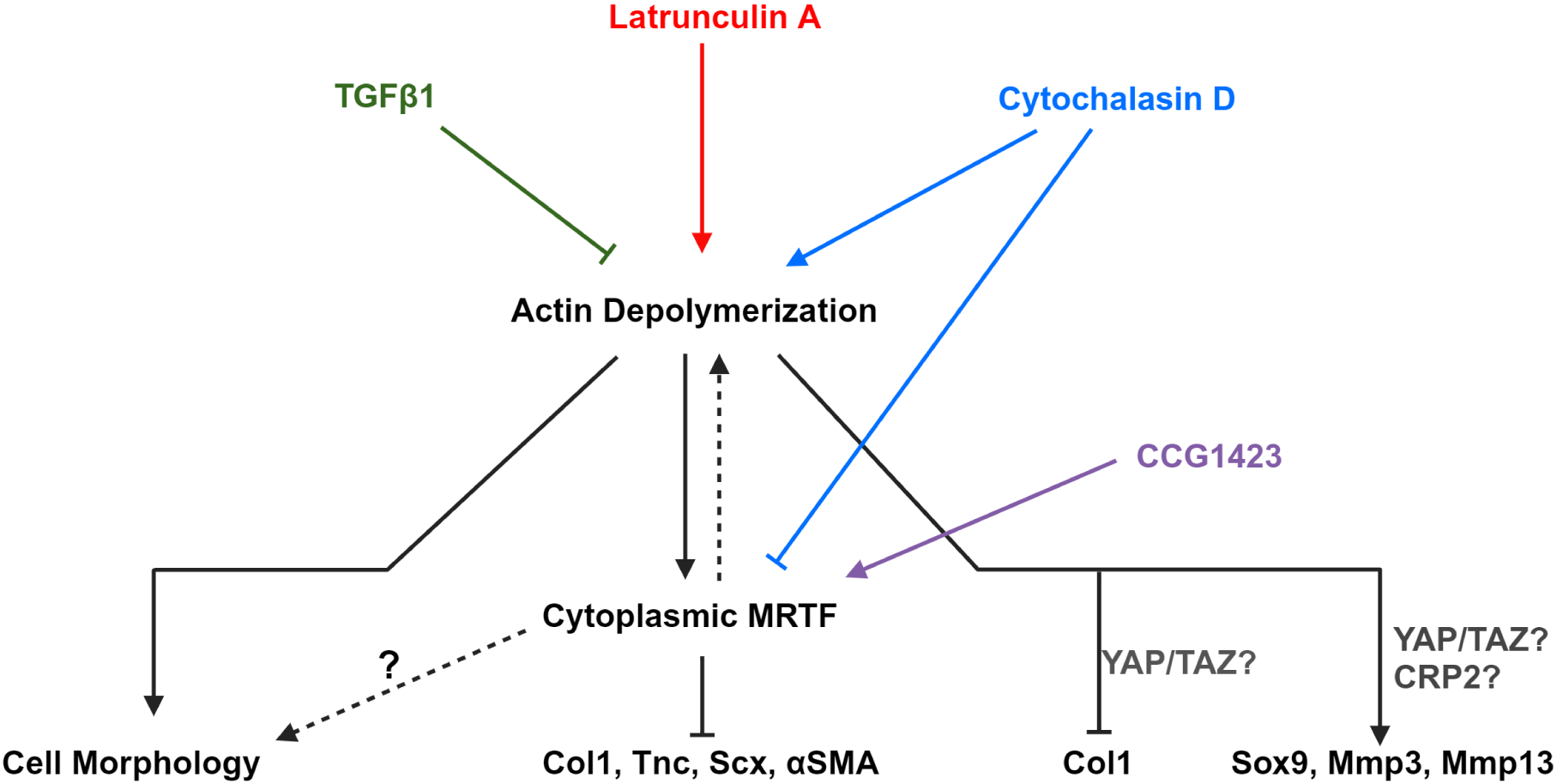
The proposed mechanism by which actin depolymerization regulates tenocyte phenotype through MRTF. Actin polymerization is a potent regulator of tenocyte phenotype through regulation of cell morphology and gene expression. Actin polymerization uses several mechanisms to regulate phenotype, including MRTF. The depolymerization of actin reduces nuclear MRTF, sequestering it in the cytoplasm, to downregulate a subset of genes associated with the tenogenic phenotype. Actin depolymerization also regulates genes in an MRTF independent fashion.

MRTF signaling alters tendon homeostasis by regulating the expression of specific molecules associated with anabolism (Figure 10). This study is the first to examine tenocyte gene regulation by actin polymerization through MRTF. The specificity of MRTF gene regulation was evident as MRTF inhibition downregulated Col1, Tnc, Scx, and αsma but did not alter Sox9, Mmp-3, and Mmp-13. The regulation of Col1, Tnc, and αsma by MRTF is consistent with previous work in other cell types (Luchsinger et al., 2011; Parreno et al., 2022; Parreno et al., 2014; Parreno et al., 2017; Yokota et al., 2017). The regulation of Scx by MRTF is a novel finding. However, Scx has previously been shown to be regulated by MRTF’s binding partner, SRF, in mesenchymal cells (Vermeulen et al., 2020).

MRTF inhibition also affected tendon cell morphology by reducing cell area and circularity and repressing stress fibers. These effects may be due to indirect/secondary effects. In support of this, MRTF inhibition had much subtler effects on tendon cells than the treatment of latrunculin A. Unlike latrunculin A, which directly affects actin, the effects of MRTF inhibition on morphology and actin could be due to indirect/secondary effects. Of the 62 genes known to be SRF target genes, 45% are cytoskeletal (Sun et al., 2006). These include molecules known to associate with F-actin and promote stress fibers, such as transgelin (Matsui, Ishikawa, & Deguchi, 2018; Shen et al., 2011), and tropomyosin (Inguito et al., 2022; Parreno, Amadeo, Kwon, & Fowler, 2020; Schofield et al., 2024; Tojkander et al., 2011). Indeed, the knockdown of SRF represses stress fiber organization in endothelial cells (Sun et al., 2006). This demonstrates a potential positive feedback loop between MRTF and F-actin depolymerization; deactivation of MRTF signaling can result in actin depolymerization, which self-propagates to cause a further reduction in nuclear MRTF (Figure 10). Considering the potent effect of actin depolymerization on causing a tendinosis-like phenotype, this could suggest that dysregulation of MRTF contributes to tendon pathology. Thus, proper MRTF signaling may be essential for maintaining tendon homeostasis and warrants future studies to delineate the role of MRTF *in vivo* in the context of tendon pathology.

F-actin polymerization status also regulates gene expression within tenocytes independent of MRTF. In keeping with previous studies (Inguito et al., 2022; Maharam et al., 2015), we showed that perturbation of F-actin networks causes a catabolic shift in the tenocyte phenotype. We determined that latrunculin A-induced depolymerization causes cell rounding, loss of F-actin stress fibers, decrease in tenogenic gene and αsma mRNA levels, as well as increases in chondrogenic (I.e., Sox9) and protease levels. Our data suggests that the regulation of Col1 may be both independent and dependent on MRTF (Figure 10), as CCG1423 and cytochalasin D led to a similar decrease in Col1 mRNA levels. The effects of cytochalasin D on Col1 contrast with what is expected if it were under a high degree of regulation by MRTF, as cytochalasin D is an MRTF activator. The effect of cytochalasin D on Col1 could be attributed to the signaling of other pathways downstream of actin polymerization, such as Yes-associated protein (YAP) and the transcriptional co-activator with PDZ-binding motif (TAZ). We previously have found support for regulating Col1 by YAP/TAZ in tenocytes (Inguito et al., 2022). We found the regulation of Sox9, Mmp3, and Mmp13 by actin depolymerization is likely independent of MRTF signaling as latrunculin and cytochalasin D had similar effects on these genes, and these genes were not affected by CCG1423 treatment. Actin depolymerization upregulates Sox9 via protein kinase A signaling (Kumar & Lassar, 2009). Furthermore, overexpression of YAP decreases Mmp3 and Mmp-13 mRNA levels (Jones et al., 2023). In addition to YAP/TAZ signaling, actin may regulate Mmp expression through an F-actin binding transcription factor, cysteine-rich protein 2 (CRP2) (Mgrditchian et al., 2023). In the case of CRP2, F-actin depolymerization liberates it from F-actin, causing translocation into the nucleus whereby it may interact with Mmp promoter regions. Thus, there are multiple pathways downstream of F-actin, aside from MRTF, which may also affect gene expression. Additional studies to investigate the contribution of other mechanisms downstream of actin require further investigation.

Actin may be at the nexus of biomechanical and biochemical signaling to regulate the tenocyte homeostasis. Previously, we determined that mechanical stress deprivation can reduce the proportion of G/F-actin in tenocytes (Inguito et al., 2022). In addition to biomechanical stimuli, in the present study, we found that exposure of tendon cells to TGFβ1 promotes F-actin polymerization and nuclear MRTF localization (Figure 10), which is consistent with other cell types (Crider et al., 2011; Gupta et al., 2013; Johnson et al., 2014; Korol et al., 2016; Kumawat et al., 2016; Speight et al., 2016). The regulation of tendon homeostasis by TGFβ has previously been attributed to canonical, Smad2/3-mediated signaling (Li et al., 2022; T. Maeda et al., 2011). Our results demonstrate that TGFβ1 also activates actin-based MRTF signaling. Thus, TGFβ may use both actin-dependent and independent pathways to regulate tenocyte homeostasis. Intriguingly, other biochemical modulators of tendon homeostasis, such as connective tissue growth factor, fibroblast growth factor-2, bone morphogenic protein, and insulin-like growth factor (Lin et al., 2023), have also been shown to regulate actin polymerization and/or MRTF signaling in other cell types (Guvakova & Surmacz, 1999; Hahn, Heusinger-Ribeiro, Lanz, Zenkel, & Goppelt-Struebe, 2000; Junglas et al., 2012; Konstantinidis, Moustakas, & Stournaras, 2011; Muehlich et al., 2007). Therefore, F-actin may be at the node of many other biochemical signals.

In conclusion, this study demonstrates that actin polymerization status regulates tenocyte expression of genes, partially through regulating the G-actin-binding transcription factor, MRTF. This work highlights that actin is a potent regulator of tendon homeostasis through regulating gene expression in tenocytes. Actin uses multiple mechanisms, including MRTF, to influence tendon homeostasis. Further understanding of the regulation of tendon homeostasis by actin and its downstream mediators during pathological processes such as tendinosis may lead to novel therapeutics to prevent disease progression and/or stimulate regeneration.

## Acknowledgments

This research was supported by the National Institutes of Health, National Institute of Arthritis and Musculoskeletal and Skin Diseases Grant R01AR080059 as well as from the Delaware Center for Musculoskeletal Research from the National Institutes of Health’s National Institute of General Medical Sciences under grant number P20GM139760. This publication was made possible by the Delaware INBRE program, supported by a grant from the National Institute of General Medical Sciences – NIGMS (P20 GM103446) from the National Institutes of Health and the State of Delaware. KLI and KMME was supported by the University of Delaware Summer Scholars Program. RM was supported by an Orthopaedic Research Society Collaborative Exchange Grant.

## Notes

### Competing Interest Statement

The authors have declared no competing interest.

## References

Arnoczky, S. P., Lavagnino, M., Egerbacher, M., Caballero, O., & Gardner, K. (2007). Matrix metalloproteinase inhibitors prevent a decrease in the mechanical properties of stress-deprived tendons: an in vitro experimental study. Am J Sports Med, 35(5), 763–769. doi:10.1177/0363546506296043

Asparuhova, M. B., Ferralli, J., Chiquet, M., & Chiquet-Ehrismann, R. (2011). The transcriptional regulator megakaryoblastic leukemia-1 mediates serum response factor-independent activation of tenascin-C transcription by mechanical stress. FASEB J, 25(10), 3477–3488. doi:10.1096/fj.11-187310

Brown, J. P., Galassi, T. V., Stoppato, M., Schiele, N. R., & Kuo, C. K. (2015). Comparative analysis of mesenchymal stem cell and embryonic tendon progenitor cell response to embryonic tendon biochemical and mechanical factors. Stem Cell Res Ther, 6(1), 89. doi:10.1186/s13287-015-0043-z

Carlier, M. F., Criquet, P., Pantaloni, D., & Korn, E. D. (1986). Interaction of cytochalasin D with actin filaments in the presence of ADP and ATP. J Biol Chem, 261(5), 2041–2050.

Chiovaro, F., Chiquet-Ehrismann, R., & Chiquet, M. (2015). Transcriptional regulation of tenascin genes. Cell Adh Migr, 9(1-2), 34–47. doi:10.1080/19336918.2015.1008333

Cooper, J. A. (1987). Effects of cytochalasin and phalloidin on actin. J Cell Biol, 105(4), 1473–1478. doi:10.1083/jcb.105.4.1473

Coue, M., Brenner, S. L., Spector, I., & Korn, E. D. (1987). Inhibition of actin polymerization by latrunculin A. FEBS Lett, 213(2), 316–318. doi:10.1016/0014-5793(87)81513-2

Crider, B. J., Risinger, G. M., Jr., Haaksma, C. J., Howard, E. W., & Tomasek, J. J. (2011). Myocardin-related transcription factors A and B are key regulators of TGF-beta1-induced fibroblast to myofibroblast differentiation. J Invest Dermatol, 131(12), 2378–2385. doi:10.1038/jid.2011.219

Dede Eren, A., Vermeulen, S., Schmitz, T. C., Foolen, J., & de Boer, J. (2023). The loop of phenotype: Dynamic reciprocity links tenocyte morphology to tendon tissue homeostasis. Acta Biomater, 163, 275–286. doi:10.1016/j.actbio.2022.05.019

Delve, E., Co, V., Regmi, S. C., Parreno, J., Schmidt, T. A., & Kandel, R. A. (2020). YAP/TAZ regulates the expression of proteoglycan 4 and tenascin C in superficial-zone chondrocytes. Eur Cell Mater, 39, 48–64. doi:10.22203/eCM.v039a03

Delve, E., Parreno, J., Co, V., Wu, P. H., Chong, J., Di Scipio, M., & Kandel, R. A. (2018). CDC42 regulates the expression of superficial zone molecules in part through the actin cytoskeleton and myocardin-related transcription factor-A. J Orthop Res, 36(9), 2421–2430. doi:10.1002/jor.23892

Egerbacher, M., Gardner, K., Caballero, O., Hlavaty, J., Schlosser, S., Arnoczky, S. P., & Lavagnino, M. (2022). Stress-deprivation induces an up-regulation of versican and connexin-43 mRNA and protein synthesis and increased ADAMTS-1 production in tendon cells in situ. Connect Tissue Res, 63(1), 43–52. doi:10.1080/03008207.2021.1873302

Espira, L., Lamoureux, L., Jones, S. C., Gerard, R. D., Dixon, I. M., & Czubryt, M. P. (2009). The basic helix-loop-helix transcription factor scleraxis regulates fibroblast collagen synthesis. J Mol Cell Cardiol, 47(2), 188–195. doi:10.1016/j.yjmcc.2009.03.024

Farhat, Y. M., Al-Maliki, A. A., Easa, A., O’Keefe, R. J., Schwarz, E. M., & Awad, H. A. (2015). TGF-beta1 Suppresses Plasmin and MMP Activity in Flexor Tendon Cells via PAI-1: Implications for Scarless Flexor Tendon Repair. J Cell Physiol, 230(2), 318–326. doi:10.1002/jcp.24707

Gardner, K., Arnoczky, S. P., Caballero, O., & Lavagnino, M. (2008). The effect of stress-deprivation and cyclic loading on the TIMP/MMP ratio in tendon cells: an in vitro experimental study. Disabil Rehabil, 30(20-22), 1523–1529. doi:10.1080/09638280701785395

Gonzalez-Nolde, S., Schweiger, C. J., Davis, E. E. R., Manzoni, T. J., Hussein, S. M. I., Schmidt, T. A., … Parreno, J. (2024). The Actin Cytoskeleton as a Regulator of Proteoglycan 4. Cartilage, 19476035231223455. doi:10.1177/19476035231223455

Gupta, M., Korol, A., & West-Mays, J. A. (2013). Nuclear translocation of myocardin-related transcription factor-A during transforming growth factor beta-induced epithelial to mesenchymal transition of lens epithelial cells. Mol Vis, 19, 1017–1028.

Guvakova, M. A., & Surmacz, E. (1999). The activated insulin-like growth factor I receptor induces depolarization in breast epithelial cells characterized by actin filament disassembly and tyrosine dephosphorylation of FAK, Cas, and paxillin. Exp Cell Res, 251(1), 244–255. doi:10.1006/excr.1999.4566

Hahn, A., Heusinger-Ribeiro, J., Lanz, T., Zenkel, S., & Goppelt-Struebe, M. (2000). Induction of connective tissue growth factor by activation of heptahelical receptors. Modulation by Rho proteins and the actin cytoskeleton. J Biol Chem, 275(48), 37429–37435. doi:10.1074/jbc.M000976200

Havis, E., Bonnin, M. A., Esteves de Lima, J., Charvet, B., Milet, C., & Duprez, D. (2016). TGFbeta and FGF promote tendon progenitor fate and act downstream of muscle contraction to regulate tendon differentiation during chick limb development. Development, 143(20), 3839–3851. doi:10.1242/dev.136242

Hayashi, K., Watanabe, B., Nakagawa, Y., Minami, S., & Morita, T. (2014). RPEL proteins are the molecular targets for CCG-1423, an inhibitor of Rho signaling. PLoS One, 9(2), e89016. doi:10.1371/journal.pone.0089016

Heinemeier, K., Langberg, H., Olesen, J. L., & Kjaer, M. (2003). Role of TGF-beta1 in relation to exercise-induced type I collagen synthesis in human tendinous tissue. J Appl Physiol (1985), 95(6), 2390–2397. doi:10.1152/japplphysiol.00403.2003

Inguito, K. L., Schofield, M. M., Faghri, A. D., Bloom, E. T., Heino, M., West, V. C., … Parreno, J. (2022). Stress deprivation of tendon explants or Tpm3.1 inhibition in tendon cells reduces F-actin to promote a tendinosis-like phenotype. Mol Biol Cell, 33(14), ar141. doi:10.1091/mbc.E22-02-0067

Ippolito, E., Natali, P. G., Postacchini, F., Accinni, L., & De Martino, C. (1977). Ultrastructural and immunochemical evidence of actin in the tendon cells. Clin Orthop Relat Res(126), 282–284.

Johnson, L. A., Rodansky, E. S., Haak, A. J., Larsen, S. D., Neubig, R. R., & Higgins, P. D. (2014). Novel Rho/MRTF/SRF inhibitors block matrix-stiffness and TGF-beta-induced fibrogenesis in human colonic myofibroblasts. Inflamm Bowel Dis, 20(1), 154–165. doi:10.1097/01.MIB.0000437615.98881.31

Jones, D. L., Hallstrom, G. F., Jiang, X., Locke, R. C., Evans, M. K., Bonnevie, E. D., Dyment, N. A. (2023). Mechanoepigenetic regulation of extracellular matrix homeostasis via Yap and Taz. Proc Natl Acad Sci U S A, 120(22), e2211947120. doi:10.1073/pnas.2211947120

Junglas, B., Kuespert, S., Seleem, A. A., Struller, T., Ullmann, S., Bosl, M., … Fuchshofer, R. (2012). Connective tissue growth factor causes glaucoma by modifying the actin cytoskeleton of the trabecular meshwork. Am J Pathol, 180(6), 2386–2403. doi:10.1016/j.ajpath.2012.02.030

Kaji, D. A., Howell, K. L., Balic, Z., Hubmacher, D., & Huang, A. H. (2020). Tgfbeta signaling is required for tenocyte recruitment and functional neonatal tendon regeneration. Elife, 9. doi:10.7554/eLife.51779

Kannus, P. (2000). Structure of the tendon connective tissue. Scand J Med Sci Sports, 10(6), 312–320. doi:10.1034/j.1600-0838.2000.010006312.x

Konstantinidis, G., Moustakas, A., & Stournaras, C. (2011). Regulation of myosin light chain function by BMP signaling controls actin cytoskeleton remodeling. Cell Physiol Biochem, 28(5), 1031–1044. doi:10.1159/000335790

Korol, A., Taiyab, A., & West-Mays, J. A. (2016). RhoA/ROCK signaling regulates TGFbeta-induced epithelial-mesenchymal transition of lens epithelial cells through MRTF-A. Mol Med, 22, 713–723. doi:10.2119/molmed.2016.00041

Kumar, D., & Lassar, A. B. (2009). The transcriptional activity of Sox9 in chondrocytes is regulated by RhoA signaling and actin polymerization. Mol Cell Biol, 29(15), 4262–4273. doi:10.1128/MCB.01779-08

Kumawat, K., Koopmans, T., Menzen, M. H., Prins, A., Smit, M., Halayko, A. J., & Gosens, R. (2016). Cooperative signaling by TGF-beta1 and WNT-11 drives sm-alpha-actin expression in smooth muscle via Rho kinase-actin-MRTF-A signaling. Am J Physiol Lung Cell Mol Physiol, 311(3), L529–537. doi:10.1152/ajplung.00387.2015

Kuo, C. K., Petersen, B. C., & Tuan, R. S. (2008). Spatiotemporal protein distribution of TGF-betas, their receptors, and extracellular matrix molecules during embryonic tendon development. Dev Dyn, 237(5), 1477–1489. doi:10.1002/dvdy.21547

Kuwahara, K., Barrientos, T., Pipes, G. C., Li, S., & Olson, E. N. (2005). Muscle-specific signaling mechanism that links actin dynamics to serum response factor. Mol Cell Biol, 25(8), 3173–3181. doi:10.1128/MCB.25.8.3173-3181.2005

Lavagnino, M., & Arnoczky, S. P. (2005). In vitro alterations in cytoskeletal tensional homeostasis control gene expression in tendon cells. J Orthop Res, 23(5), 1211–1218. doi:10.1016/j.orthres.2005.04.001

Lejard, V., Brideau, G., Blais, F., Salingcarnboriboon, R., Wagner, G., Roehrl, M. H., … Rossert, J. (2007). Scleraxis and NFATc regulate the expression of the pro-alpha1(I) collagen gene in tendon fibroblasts. J Biol Chem, 282(24), 17665–17675. doi:10.1074/jbc.M610113200

Li, Y., Liu, X., Liu, X., Peng, Y., Zhu, B., Guo, S., … Li, S. (2022). Transforming growth factor-beta signalling pathway in tendon healing. Growth Factors, 40(3-4), 98–107. doi:10.1080/08977194.2022.2082294

Lin, M., Li, W., Ni, X., Sui, Y., Li, H., Chen, X., Wang, C. (2023). Growth factors in the treatment of Achilles tendon injury. Front Bioeng Biotechnol, 11, 1250533. doi:10.3389/fbioe.2023.1250533

Luchsinger, L. L., Patenaude, C. A., Smith, B. D., & Layne, M. D. (2011). Myocardin-related transcription factor-A complexes activate type I collagen expression in lung fibroblasts. J Biol Chem, 286(51), 44116–44125. doi:10.1074/jbc.M111.276931

Maeda, E., Sugimoto, M., & Ohashi, T. (2013). Cytoskeletal tension modulates MMP-1 gene expression from tenocytes on micropillar substrates. J Biomech, 46(5), 991–997. doi:10.1016/j.jbiomech.2012.11.056

Maeda, T., Sakabe, T., Sunaga, A., Sakai, K., Rivera, A. L., Keene, D. R., Sakai, T. (2011). Conversion of mechanical force into TGF-beta-mediated biochemical signals. Curr Biol, 21(11), 933–941. doi:10.1016/j.cub.2011.04.007

Maharam, E., Yaport, M., Villanueva, N. L., Akinyibi, T., Laudier, D., He, Z., Sun, H. B. (2015). Rho/Rock signal transduction pathway is required for MSC tenogenic differentiation. Bone Res, 3, 15015. doi:10.1038/boneres.2015.15

Matsui, T. S., Ishikawa, A., & Deguchi, S. (2018). Transgelin-1 (SM22alpha) interacts with actin stress fibers and podosomes in smooth muscle cells without using its actin binding site. Biochem Biophys Res Commun, 505(3), 879–884. doi:10.1016/j.bbrc.2018.09.176

Medjkane, S., Perez-Sanchez, C., Gaggioli, C., Sahai, E., & Treisman, R. (2009). Myocardin-related transcription factors and SRF are required for cytoskeletal dynamics and experimental metastasis. Nat Cell Biol, 11(3), 257–268. doi:10.1038/ncb1833

Mehr, D., Pardubsky, P. D., Martin, J. A., & Buckwalter, J. A. (2000). Tenascin-C in tendon regions subjected to compression. J Orthop Res, 18(4), 537–545. doi:10.1002/jor.1100180405

Mgrditchian, T., Brown-Clay, J., Hoffmann, C., Muller, T., Filali, L., Ockfen, E., Thomas, C. (2023). Actin cytoskeleton depolymerization increases matrix metalloproteinase gene expression in breast cancer cells by promoting translocation of cysteine-rich protein 2 to the nucleus. Front Cell Dev Biol, 11, 1100938. doi:10.3389/fcell.2023.1100938

Miralles, F., Posern, G., Zaromytidou, A. I., & Treisman, R. (2003). Actin dynamics control SRF activity by regulation of its co-activator MAL. Cell, 113(3), 329–342. doi:10.1016/s0092-8674(03)00278-2

Motulsky, H. J., & Brown, R. E. (2006). Detecting outliers when fitting data with nonlinear regression - a new method based on robust nonlinear regression and the false discovery rate. BMC Bioinformatics, 7, 123. doi:10.1186/1471-2105-7-123

Mouilleron, S., Langer, C. A., Guettler, S., McDonald, N. Q., & Treisman, R. (2011). Structure of a pentavalent G-actin*MRTF-A complex reveals how G-actin controls nucleocytoplasmic shuttling of a transcriptional co-activator. Sci Signal, 4(177), ra40. doi:10.1126/scisignal.2001750

Mousavizadeh, R., West, V. C., Inguito, K. L., Elliott, D. M., & Parreno, J. (2023). The application of mechanical load onto mouse tendons by magnetic restraining represses Mmp-3 expression. BMC Res Notes, 16(1), 127. doi:10.1186/s13104-023-06413-z

Muehlich, S., Cicha, I., Garlichs, C. D., Krueger, B., Posern, G., & Goppelt-Struebe, M. (2007). Actin-dependent regulation of connective tissue growth factor. Am J Physiol Cell Physiol, 292(5), C1732–1738. doi:10.1152/ajpcell.00552.2006

Nalluri, S. M., O’Connor, J. W., & Gomez, E. W. (2015). Cytoskeletal signaling in TGFbeta-induced epithelial-mesenchymal transition. Cytoskeleton (Hoboken), 72(11), 557–569. doi:10.1002/cm.21263

Parreno, J., Amadeo, M. B., Kwon, E. H., & Fowler, V. M. (2020). Tropomyosin 3.1 Association With Actin Stress Fibers is Required for Lens Epithelial to Mesenchymal Transition. Invest Ophthalmol Vis Sci, 61(6), 2. doi:10.1167/iovs.61.6.2

Parreno, J., Emin, G., Vu, M. P., Clark, J. T., Aryal, S., Patel, S. D., & Cheng, C. (2022). Methodologies to unlock the molecular expression and cellular structure of ocular lens epithelial cells. Front Cell Dev Biol, 10, 983178. doi:10.3389/fcell.2022.983178

Parreno, J., Raju, S., Niaki, M. N., Andrejevic, K., Jiang, A., Delve, E., & Kandel, R. (2014). Expression of type I collagen and tenascin C is regulated by actin polymerization through MRTF in dedifferentiated chondrocytes. FEBS Lett, 588(20), 3677–3684. doi:10.1016/j.febslet.2014.08.012

Parreno, J., Raju, S., Wu, P. H., & Kandel, R. A. (2017). MRTF-A signaling regulates the acquisition of the contractile phenotype in dedifferentiated chondrocytes. Matrix Biol, 62, 3–14. doi:10.1016/j.matbio.2016.10.004

Pryce, B. A., Watson, S. S., Murchison, N. D., Staverosky, J. A., Dunker, N., & Schweitzer, R. (2009). Recruitment and maintenance of tendon progenitors by TGFbeta signaling are essential for tendon formation. Development, 136(8), 1351–1361. doi:10.1242/dev.027342

Ralphs, J. R., Waggett, A. D., & Benjamin, M. (2002). Actin stress fibres and cell-cell adhesion molecules in tendons: organisation in vivo and response to mechanical loading of tendon cells in vitro. Matrix Biol, 21(1), 67–74. doi:10.1016/s0945-053x(01)00179-2

Schmittgen, T. D., & Livak, K. J. (2008). Analyzing real-time PCR data by the comparative C(T) method. Nat Protoc, 3(6), 1101–1108. doi:10.1038/nprot.2008.73

Schofield, M. M., Rzepski, A. T., Richardson-Solorzano, S., Hammerstedt, J., Shah, S., Mirack, C. E., … Parreno, J. (2024). Targeting F-actin stress fibers to suppress the dedifferentiated phenotype in chondrocytes. Eur J Cell Biol, 103(2), 151424. doi:10.1016/j.ejcb.2024.151424

Screen, H. R., Berk, D. E., Kadler, K. E., Ramirez, F., & Young, M. F. (2015). Tendon functional extracellular matrix. J Orthop Res, 33(6), 793–799. doi:10.1002/jor.22818

Shen, J., Yang, M., Jiang, H., Ju, D., Zheng, J. P., Xu, Z., … Li, L. (2011). Arterial injury promotes medial chondrogenesis in Sm22 knockout mice. Cardiovasc Res, 90(1), 28–37. doi:10.1093/cvr/cvq378

Spector, I., Shochet, N. R., Kashman, Y., & Groweiss, A. (1983). Latrunculins: novel marine toxins that disrupt microfilament organization in cultured cells. Science, 219(4584), 493–495. doi:10.1126/science.6681676

Spector, M. (2001). Musculoskeletal connective tissue cells with muscle: expression of muscle actin in and contraction of fibroblasts, chondrocytes, and osteoblasts. Wound Repair Regen, 9(1), 11–18. doi:10.1046/j.1524-475x.2001.00011.x

Speight, P., Kofler, M., Szaszi, K., & Kapus, A. (2016). Context-dependent switch in chemo/mechanotransduction via multilevel crosstalk among cytoskeleton-regulated MRTF and TAZ and TGFbeta-regulated Smad3. Nat Commun, 7, 11642. doi:10.1038/ncomms11642

Sun, Q., Chen, G., Streb, J. W., Long, X., Yang, Y., Stoeckert, C. J., Jr., & Miano, J. M. (2006). Defining the mammalian CArGome. Genome Res, 16(2), 197–207. doi:10.1101/gr.4108706

Tojkander, S., Gateva, G., Schevzov, G., Hotulainen, P., Naumanen, P., Martin, C., … Lappalainen, P. (2011). A molecular pathway for myosin II recruitment to stress fibers. Curr Biol, 21(7), 539–550. doi:10.1016/j.cub.2011.03.007

Vermeulen, S., Roumans, N., Honig, F., Carlier, A., Hebels, D., Eren, A. D., … de Boer, J. (2020). Mechanotransduction is a context-dependent activator of TGF-beta signaling in mesenchymal stem cells. Biomaterials, 259, 120331. doi:10.1016/j.biomaterials.2020.120331

Wunderli, S. L., Blache, U., Beretta Piccoli, A., Niederost, B., Holenstein, C. N., Passini, F. S., … Snedeker, J. G. (2020). Tendon response to matrix unloading is determined by the patho-physiological niche. Matrix Biol, 89, 11–26. doi:10.1016/j.matbio.2019.12.003

Xie, J., Ning, Q., Zhang, H., Ni, S., & Ye, J. (2022). RhoA/ROCK Signaling Regulates TGF-beta1-Induced Fibrotic Effects in Human Pterygium Fibroblasts through MRTF-A. Curr Eye Res, 47(2), 196–205. doi:10.1080/02713683.2021.1962363

Yokota, S., Chosa, N., Kyakumoto, S., Kimura, H., Ibi, M., Kamo, M., … Ishisaki, A. (2017). ROCK/actin/MRTF signaling promotes the fibrogenic phenotype of fibroblast-like synoviocytes derived from the temporomandibular joint. Int J Mol Med, 39(4), 799–808. doi:10.3892/ijmm.2017.2896

